# DDI_single: Single-Sequence-Based Protein Domain Assembly

**DOI:** 10.64898/2026.06.05.730531

**Authors:** Zong Shengyi

## Abstract

Domains are the basic units of protein structure and function. Appropriate inter-domain organization is critical to enable cooperative execution of multiple related functions. It is thus a crucial step to determine the full-length structure of multi-domain proteins for the purpose of elucidating their functions and designing new drugs to regulate these functions.

Existing structure prediction algorithms are generally better at solving the internal conformation of domains, rather than modeling the relative positions between domains. To address the challenge of accurately determining multi-domain protein conformations, we develop a single-sequence-based domain assembly algorithm called DDI_single. DDI_single directly extracts features from the amino acid sequence using the protein language model ESM-lb, and accurately predicts the interactions between residue pairs of structural domains through a novel gated cross-attention module, thus achieving the correct assembly of structural domains.

With the knowledge of domain definition, DDI_single achieves more than 20% higher accuracy in the task of predicting the relative distances of residue pairs between domains than that of the single-sequence-based structure prediction algorithm trRosettaX_single. When assembling domains with known spatial conformations, DDI_single correctly assembles 74.4% of the samples in the test set (TM-score*>*0.5). When assembling domains with unknown spatial conformations, in cases where the internal spatial conformations of domains are correctly modeled, DDI_single correctly assembles 73.9% of the samples.

## 1 Introduction

The spatial structure of a protein determines its biological function. While experimental methods such as X-ray crystallography [1], NMR spectroscopy [2], and cryo-EM [3] can resolve structures, they remain costly and time-consuming. As of 2023, the Protein Data Bank (PDB) [4] contains only ∼200,000 entries—less than one-thousandth of known protein sequences [5]. Computational structure prediction methods, including homology modeling [6], *ab initio* prediction [7], and deep learning approaches such as trRosettaX [8] and AlphaFold2 [10], have made remarkable progress but still face challenges for multi-domain proteins.

At least 80% of eukaryotic proteins and 67% of prokaryotic proteins are composed of two or more structural domains [17]. The proper spatial arrangement of domains is critical for cooperative function execution, and active sites of multi-domain enzymes are often located at domain interfaces. However, existing prediction methods are generally better at modeling intra-domain conformations than inter-domain relative positions, because template-based methods struggle to find templates spanning multiple domains, and self-attention mechanisms in single-sequence methods tend to underweight the weaker inter-domain interactions relative to stronger intra-domain ones.

Single-sequence methods based on protein language models (PLMs), such as trRosettaX_single [16], bypass the need for homologous sequences by extracting features directly from amino acid sequences using PLMs like ESM-1b [13]. This eliminates the costly MSA search step and enables prediction for orphan proteins. However, these methods still underperform on inter-domain distance prediction due to the self-attention limitation described above.

To address this, we propose DDI_single, a single-sequence-based domain assembly algorithm that uses a novel gated cross-attention module to specifically capture inter-domain residue pair interactions. Given domain boundary information, DDI_single splits the PLM embedding by domain, applies cross-attention between domains with an expanded query/key space and a gating mechanism inspired by AlphaFold2 and LSTM networks [23], and predicts inter-domain distances and orientations (*d, ω, θ, φ*). The predictions are integrated with intra-domain constraints and assembled via Rosetta energy minimization.

Our main contributions are:

- A gated cross-attention module with expanded query/key dimensions that focuses specifically on inter-domain residue pair interactions.
- A weight-sharing recycling mechanism that increases network depth without proportional parameter growth.
- DDI_single achieves *>*20% higher inter-domain distance prediction accuracy than trRoset-taX_single, correctly assembling 74.4% of test proteins with known domain conformations (TM-score*>*0.5) and 73.9% when domain conformations are predicted.

## 2 Related Work

### Structure Prediction

Mainstream methods rely on homologous sequences and structural templates. trRosettaX [8] combines deep learning with template-based approaches, achieving top performance at CASP15 [9]. AlphaFold2 [10] uses an end-to-end architecture with Evoformer to achieve near-experimental accuracy. However, both require expensive MSA searches and struggle when homologous sequences are scarce [11].

### Single-Sequence Methods

Protein language models such as ESM-1b [13] and ESM-2 [14], trained via masked language modeling on large sequence databases [15], encode rich structural information from sequences alone. trRosettaX_single [16] leverages ESM-1b to predict inter-residue distances and orientations without MSAs, achieving strong results on orphan proteins.

### Domain Assembly

Multi-domain proteins comprise at least 80% of eukaryotic proteomes [17], yet predicting inter-domain arrangements remains challenging. Template-based methods rarely find multi-domain templates, and single-sequence methods using global self-attention tend to under-weight weaker inter-domain interactions. Our work specifically targets this gap through a cross-attention mechanism focused on inter-domain residue pairs.

## 3 Method

Figure 1 illustrates the DDI_single pipeline. The model encodes the input sequence via ESM-1b, splits the embedding by domain boundaries, applies a gated cross-attention-based feature extraction module with weight-sharing recycling [10] (4 iterations), predicts inter-domain distances and orientations (*d, ω, θ, φ*), and assembles domains via Rosetta energy minimization. For proteins with three or more domains, pairs are processed sequentially.

**Figure 1:**
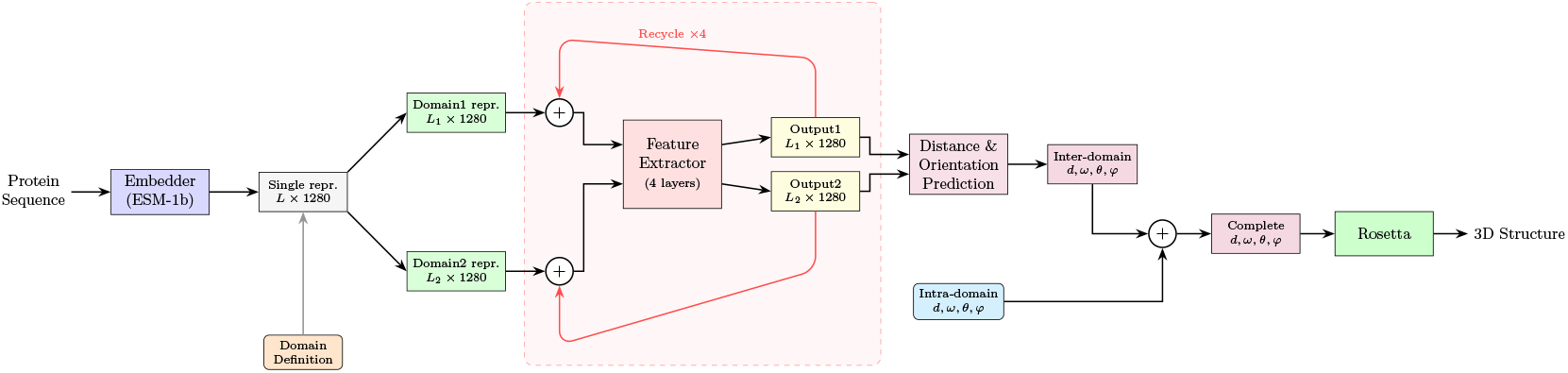
The overall pipeline of DDI_single. The model takes a protein sequence as input, extracts features using ESM-1b, splits representations by domain boundaries, processes them through the gated cross-attention-based feature extraction module with recycling, predicts inter-domain distances and orientations (*d, ω, θ, φ*), and assembles domains via Rosetta energy minimization.

### 3.1 Feature Embedding Module

We use the pre-trained protein language model ESM-1b [13] (33 transformer layers, output dimension 1280) as the feature embedding module without architectural modification.

### 3.2 Feature Extraction Module

The feature extraction module (Figure 2) consists of a single unit containing a gated cross-attention layer followed by a feedforward layer with residual connections [19, 20]. This unit is applied iteratively 4 times with shared weights (recycling) [10]: the output representations are fed back as input for the next iteration, progressively refining the inter-domain features without increasing parameter count. The two domain representations are updated in parallel through inter-domain cross-attention at each iteration.

**Figure 2:**
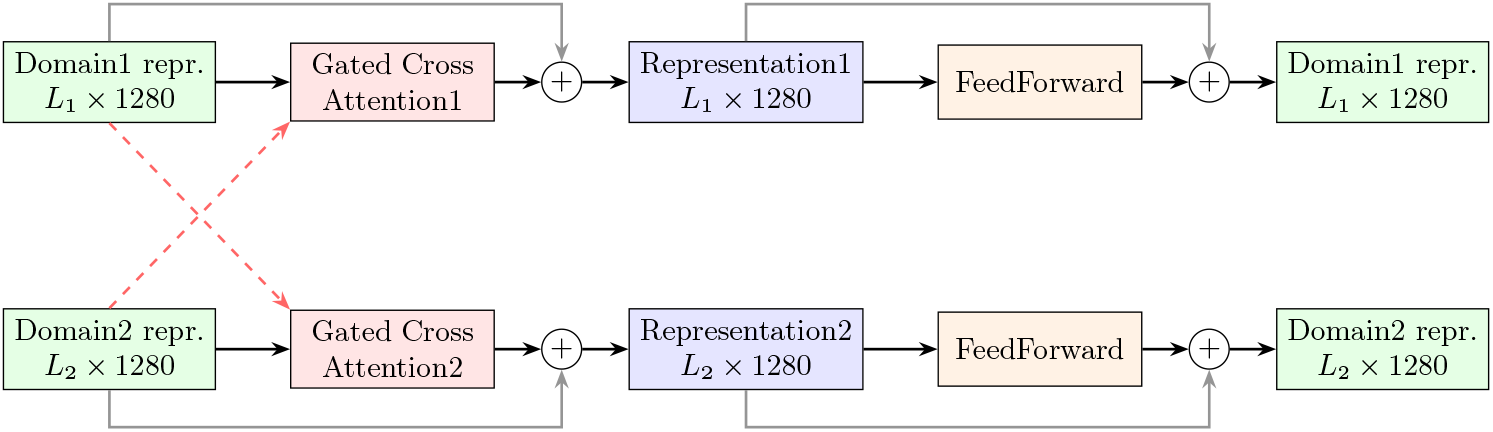
Architecture of the DDI_single feature extraction module. Domain 1 and Domain 2 representations are processed through gated cross-attention layers followed by feedforward layers with residual connections.

#### 3.2.1 Gated Cross-Attention Layer

We design a gated cross-attention layer (Figure 3) with three key innovations over standard self-attention: (1) **Cross-attention:** rather than computing global self-attention where inter-domain interactions are diluted, we use cross-attention [21] where one domain’s features are updated under the guidance of the other domain; (2) **Expanded query/key space:** we expand *Q* and *K* dimensions to 4*d* while keeping *V* at *d*, enlarging the interaction space [22] without increasing output dimensionality; (3) **Gating:** inspired by AlphaFold2 and LSTM [23], an independent gating branch applies Sigmoid-gated filtering to the attention output, allowing the model to dynamically enhance or suppress features.

**Figure 3:**
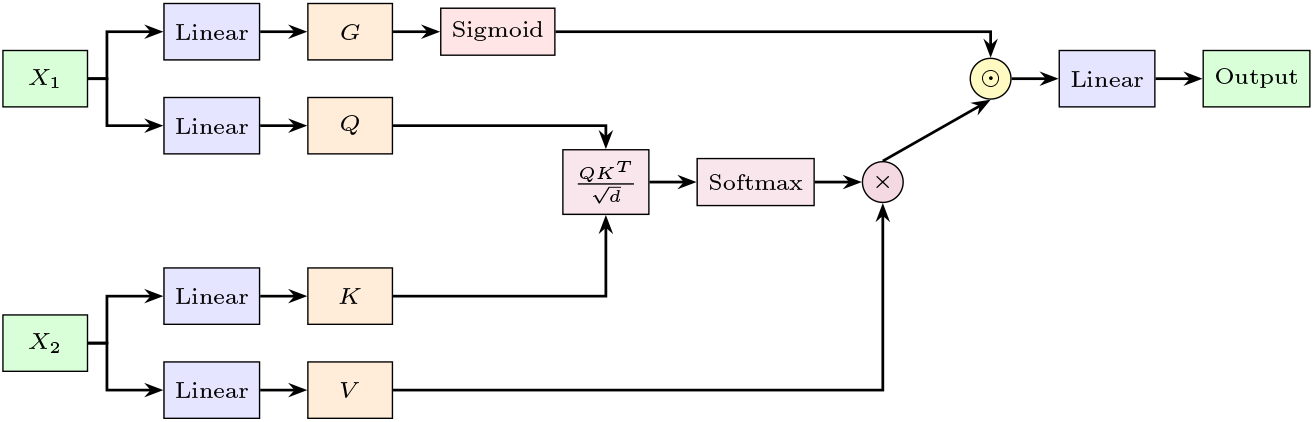
Gated cross-attention mechanism. *X*_1_ is projected into gate *G* and query *Q* (expanded to 4*d*); *X*_2_ is projected into key *K* (4*d*) and value *V* (*d*). The attention output is element-wise gated by Sigmoid(*G*).

Each gated cross-attention layer is followed by a feedforward layer with GELU [24] activation and residual connections [19].

### 3.3 Distance and Orientation Prediction Module

The distance and orientation prediction module (Figure 4) predicts the relative distances and orientations between residue pairs of structural domains based on the feature vectors of Domain 1 and Domain 2 extracted by the feature extraction module. Specifically, for each inter-domain residue pair, the module predicts four geometric quantities: the inter-residue *C*_*β*_−*C*_*β*_ distance *d*, the dihedral angle *ω* describing the rotation of both *C*_*α*_ atoms about the *C*_*β*_−*C*_*β*_ axis, the dihedral angle *θ* describing the rotation of the first residue’s nitrogen and the second residue’s *C*_*α*_ about the first residue’s *C*_*α*_−*C*_*β*_ axis, and the planar angle *φ* formed by *C*_*α,i*_−*C*_*β,i*_−*C*_*β,j*_. The distance *d* is discretized into 37 bins, while the angles *ω, θ*, and *φ* are discretized into 25, 25, and 13 bins respectively. Feature vectors from both domains are projected through linear layers, combined via outer product to form an inter-domain feature map, and then fed into four parallel branches, each containing an independent convolutional neural network (CNN), to predict the probability distributions of *d, ω, θ*, and *φ* respectively.

**Figure 4:**
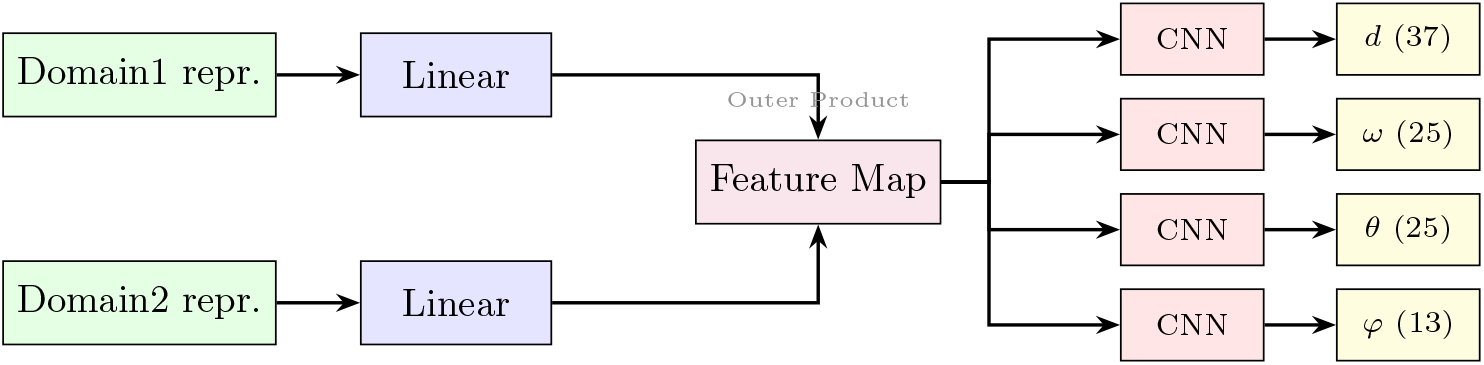
Architecture of the distance and orientation prediction module. Feature vectors from both domains are projected through linear layers, combined via outer product to form a feature map, and processed by a CNN with four output heads to predict distance (*d*, 37 bins) and orientation (*ω*, 25 bins; *θ*, 25 bins; *φ*, 13 bins) distributions.

### 3.4 Energy Minimization Modeling

Our energy minimization modeling approach follows that of trRosettaX_single. The predicted inter-residue relative distances and orientation angles (*d, ω, θ, φ*) are first converted to energy functions as spatial constraints, and then Rosetta’s quasi-Newton method is used to optimize the degrees of freedom of the model, yielding 120 coarse-grained centroid models. The top 5 lowest-energy models are selected for full-atom optimization, and the lowest-energy full-atom model is chosen as the final modeling result.

### 3.5 Training

We employ a two-step fine-tuning strategy with multi-task training [25]. Step 1 freezes the ESM-1b parameters and trains the feature extraction and prediction modules using cross-entropy loss, simultaneously predicting both inter-domain and intra-domain distances/orientations (with 2× weight on inter-domain loss). Step 2 unfreezes all parameters and adds masked language modeling [12] as an auxiliary task. We use Adam [26] with learning rate 10^−5^, warmup [27], dropout [29] (0.3), dynamic masking [28], and L2 regularization (10^−4^). Full training details are in Appendix C.

## 4 Experiments

### 4.1 Datasets

We collected multi-domain protein structures from PDB using CATH [18] domain annotations. Training data uses structures released before May 1, 2018 (same cutoff as trRosettaX_single for fair comparison); the test set uses structures from May 2018 to March 2023. Sequences are clustered at 40% similarity. For each residue pair, we compute *C*_*β*_ distances and orientation angles (*d, ω, θ, φ*), discretized into 37, 25, 25, and 13 bins respectively. Table 1 summarizes the datasets. Full preprocessing details are in Appendix B.

**Table 1:** Summary of datasets.

| Dataset | Date Range | Size |
| --- | --- | --- |
| Training set | Before May 1, 2018 | 4,524 classes |
| Evaluation set | Before May 1, 2018 | 500 classes |
| Test set | May 1, 2018 – March 29, 2023 | 681 classes |

### 4.2 Evaluation Metrics

We use **Precision** to evaluate distance prediction quality: among the top-*N* residue pairs predicted to be in contact (2−8 Å), we measure the fraction that are true contacts. We report Top10, Top5, and Top1 for inter-domain pairs, and Top*L* and Top*L*/2 for full-length sequences. We use **TM-score** [30] to evaluate 3D assembly quality; TM-score*>*0.5 indicates correct fold. Formal definitions are in Appendix D.

### 4.3 Assembly with Known Domain Conformations

In the scenario where domain conformations are known, we first extract internal residue pair relative distances and orientations from structure files, then use DDI_single and trRosettaX_single to predict inter-domain residue pair relative distances and orientations respectively, concatenate intra-domain and inter-domain constraints to obtain full-length distance and orientation matrices, and finally use Rosetta to assemble domains under both sets of spatial constraints.

#### 4.3.1 Inter-domain Distance Prediction

Table 2 compares DDI_single and trRosettaX_single on inter-domain relative distance prediction. Neither method uses ensemble prediction. Under known domain partitioning, DDI_single achieves higher prediction accuracy at all levels than trRosettaX_single, demonstrating DDI_single’s ability to accurately capture inter-domain residue pair interactions and its potential for domain assembly tasks.

**Table 2:**
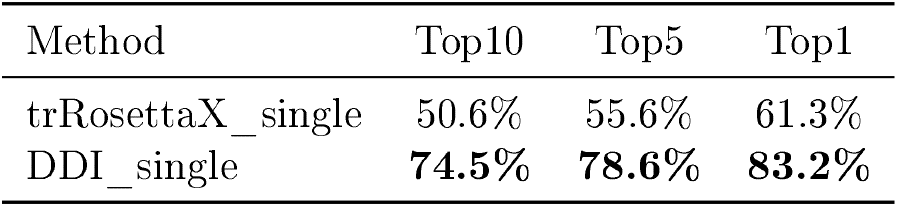
Precision of DDI_single and trRosettaX_single on inter-domain distance prediction.

#### 4.3.2 Domain Assembly

We use Rosetta to model based on the distance matrices assembled from DDI_single and trRoset-taX_single predictions respectively, using TM-score to measure domain assembly quality. Since intra-domain spatial constraints are directly extracted from protein structure files, we can consider the TM-score metric to be related only to domain assembly quality.

DDI_single assembles domains with significantly higher quality than trRosettaX_single. Among the 681 test samples, 481 samples modeled using DDI_single predictions as constraints have higher quality than those using trRosettaX_single (TM-score difference *>* 0.02), 138 samples have similar quality (absolute TM-score difference *<* 0.02), and only 62 samples show lower quality with DDI_single.

DDI_single predictions provide more accurate spatial constraints for the folding algorithm, enabling precise domain assembly. In the test set of 681 samples, DDI_single constraints help correctly assemble 507 proteins (TM-score*>*0.5), achieving a 74.4% accuracy rate with a mean TM-score of 0.7332. In contrast, trRosettaX_single’s weaker ability to extract inter-domain interactions results in poorer domain assembly quality; trRosettaX_single constraints correctly assemble only 303 proteins with 44.5% accuracy and a mean TM-score of only 0.4974. Figure 5 shows the comparison of TM-scores between the two methods. Detailed case studies are provided in Appendix A.

**Figure 5:**
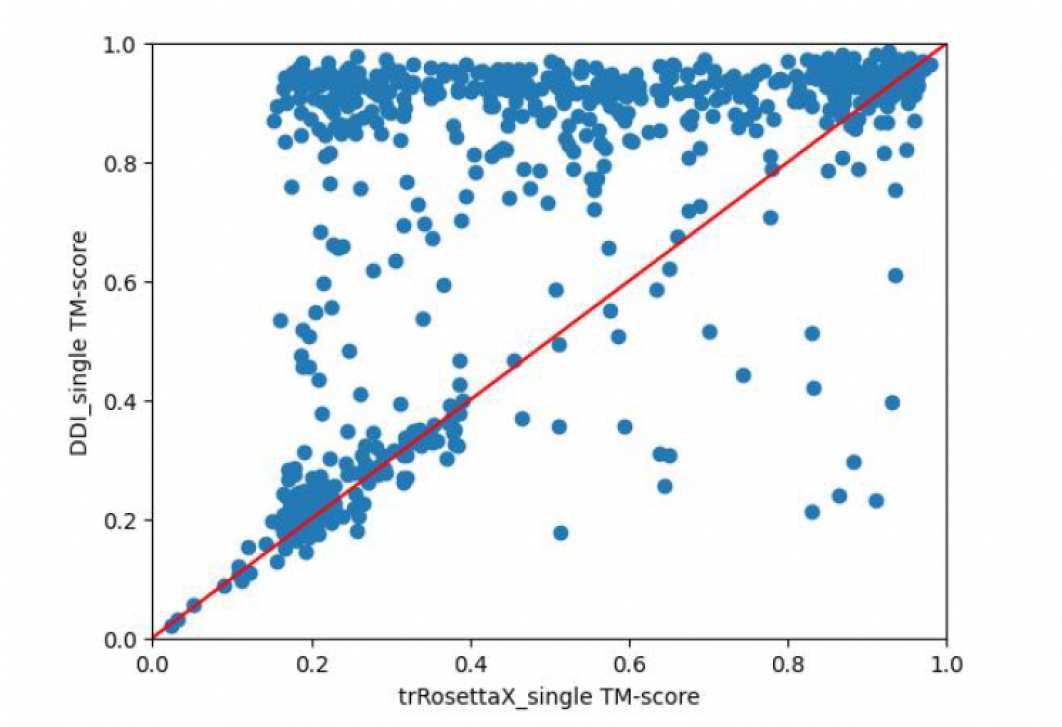
Comparison of TM-scores for domain assembly with known domain conformations. Each point represents a protein in the test set; points above the diagonal indicate DDI_single achieves higher assembly quality.

### 4.4 Assembly with Unknown Domain Conformations

Since most structural domains do not have experimentally determined conformations, evaluating DDI_single’s ability to assemble predicted domains is particularly important. In this scenario, we first use trRosettaX_single to predict the complete distance and orientation matrices of multi-domain proteins, then replace the inter-domain residue pair relative distances and orientations with DDI_single predictions, and finally use Rosetta to model and assemble the predicted structural domains.

#### 4.4.1 Full-Length Sequence Distance Prediction

Table 3 compares the full-sequence distance prediction precision before and after substitution with DDI_single. Although DDI_single significantly improves inter-domain distance and orientation prediction accuracy, because the number of inter-domain residue pairs with distance less than 8 Å is relatively small, the substitution does not bring a dramatic improvement in overall sequence precision.

**Table 3:** Precision of the full-length distance matrix before and after replacing inter-domain distances with DDI_single predictions.

| Method | TopL | TopL/2 |
| --- | --- | --- |
| trRosettaX_single | 73.1% | 82.0% |
| DDI_single | **74.1%** | **82.6%** |

#### 4.4.2 Domain Assembly

The majority of samples modeled using DDI_single have higher quality than those using trRoset-taX_single (Figure 6). In the test set of 681 samples, trRosettaX_single’s direct modeling mean TM-score is 0.3203, while using DDI_single’s predicted spatial constraints to replace trRoset-taX_single predictions gives a mean TM-score of 0.3890. Although DDI_single achieves higher domain assembly quality, both methods produce relatively poor overall folding quality with mean TM-scores below 0.5.

**Figure 6:**
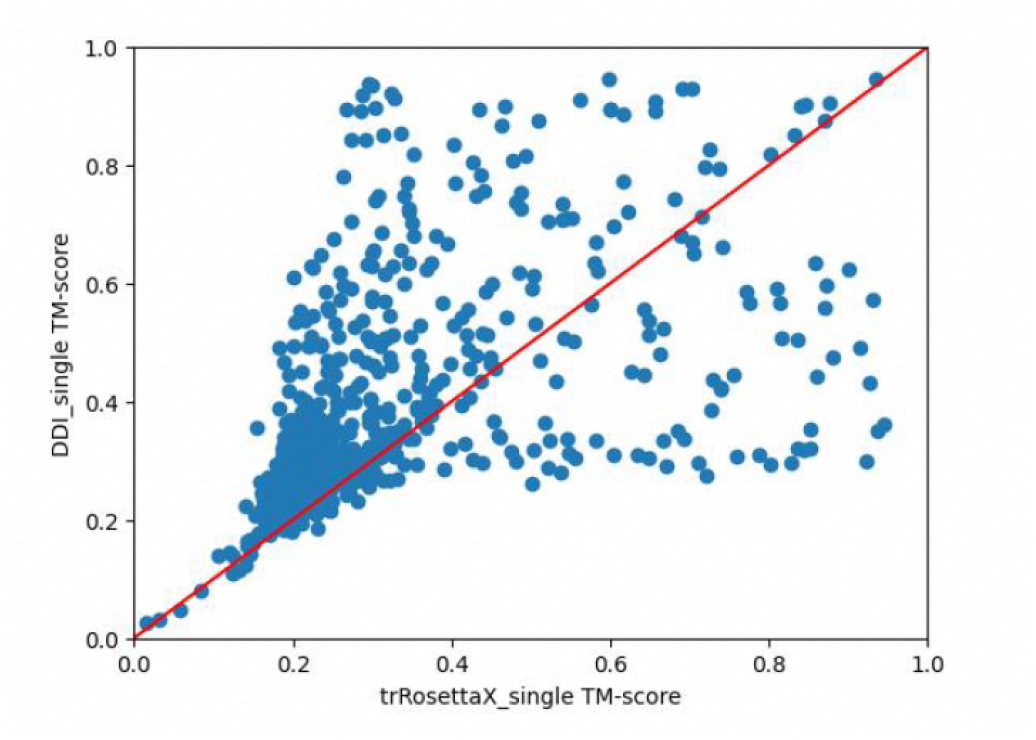
Comparison of TM-scores for domain assembly with unknown domain conformations on the full test set (681 proteins).

To investigate, we evaluated both methods’ modeling of individual domains, finding mean single-domain TM-scores of only 0.3806 (trRosettaX_single) and 0.4154 (DDI_single), well below the ideal threshold (TM-score*>*0.5), indicating that evaluation metrics are affected by incorrect intra-domain conformations.

To reduce the impact of poor intra-domain conformations on domain assembly evaluation metrics, we selected proteins where the modeled intra-domain TM-score exceeds 0.5 as a new test set and re-evaluated both methods’ domain assembly capabilities.

In the filtered subset of 115 proteins, DDI_single and trRosettaX_single model individual do-mains with mean TM-scores of 0.6745 and 0.6566 respectively, while their complete structure mean TM-scores are 0.6324 and 0.5698. For similar-quality individual domains, DDI_single’s predicted results can more accurately assemble single domains into complete structures, achieving a mean TM-score 11.0% higher than trRosettaX_single alone.

Furthermore, DDI_single correctly assembles 85 of the 115 proteins (TM-score*>*0.5), achieving an assembly accuracy of 73.9%, while trRosettaX_single can only correctly assemble 67 proteins with 58.3% accuracy (Figure 7).

**Figure 7:**
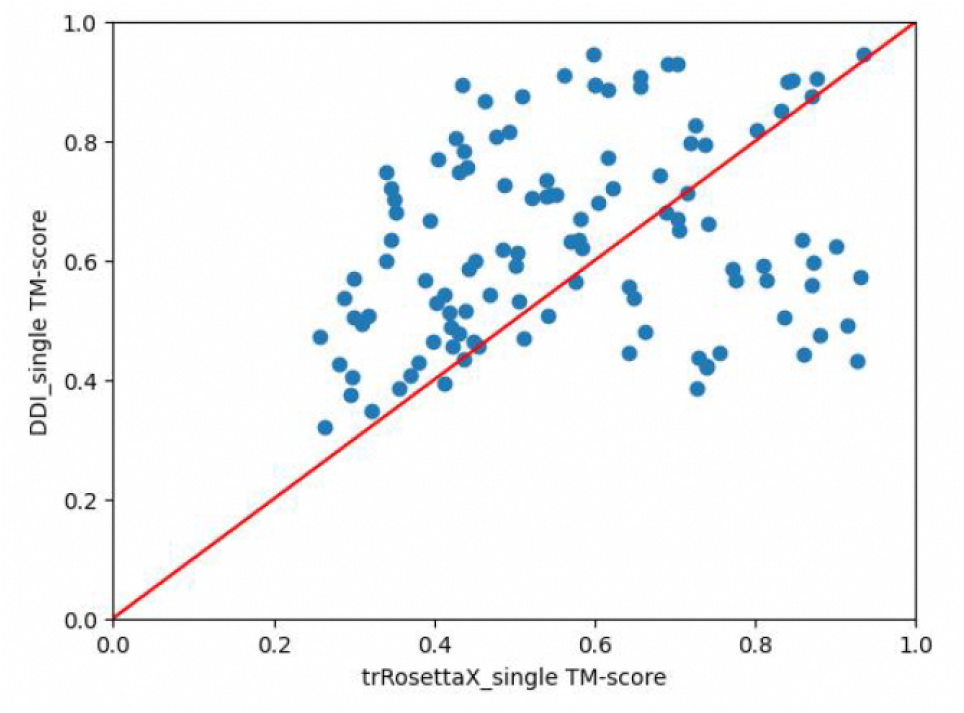
Comparison of TM-scores for domain assembly with unknown domain conformations on the filtered subset (115 proteins where intra-domain modeling TM-score *>* 0.5).

## 5 Ablation Study

To further analyze and evaluate the contributions of each innovative improvement to overall model performance, we conducted ablation experiments by systematically removing each improvement and quantifying its impact on model performance. We trained 5 ablation models using identical hyperparameters:

1. **Model 1**: Uses representative sequences from each class for training; uses traditional multi-head self-attention for feature extraction; trains only the feature extraction and distance/orientation prediction modules; does not use multi-task training; no additional training techniques.
2. **Model 2:** Uses representative sequences; uses the improved gated cross-attention module for feature extraction; trains only the feature extraction and distance/orientation prediction modules; no multi-task training; no additional training techniques.
3. **Model 3:** Randomly samples training data from each class; uses gated cross-attention; trains feature extraction and distance/orientation prediction modules; no multi-task training; uses additional training techniques to improve performance.
4. **Model 4:** Random sampling; gated cross-attention; trains feature extraction and distance/orientation prediction modules; uses multi-task training; uses additional training techniques.
5. **Model 5:** Random sampling; gated cross-attention; uses weight-sharing recycling to increase feature extraction module depth; trains feature extraction and distance/orientation prediction modules; uses multi-task training; uses additional training techniques.

Table 4 compares the prediction precision of each ablation model with DDI_single on the independent test set.

**Table 4:**
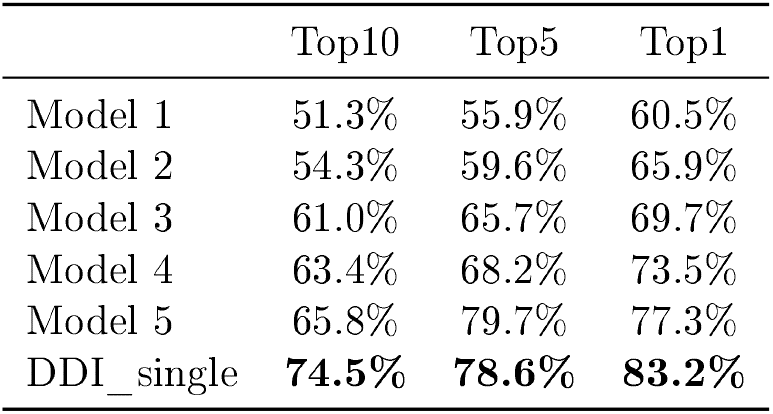
Prediction precision of ablation models and DDI_single on the test set.

|  | Top10 | Top5 | Top1 |
| --- | --- | --- | --- |
| Model 1 | 51.3% | 55.9% | 60.5% |
| Model 2 | 54.3% | 59.6% | 65.9% |
| Model 3 | 61.0% | 65.7% | 69.7% |
| Model 4 | 63.4% | 68.2% | 73.5% |
| Model 5 | 65.8% | 79.7% | 77.3% |
| DDI_single | **74.5%** | **78.6%** | **83.2%** |

From the table, several conclusions can be drawn: (1) The improved gated cross-attention module forces the model to focus on inter-domain residue pair interactions, with Model 2’s prediction accuracy far exceeding the baseline Model 1 using traditional multi-head self-attention. (2) Increasing training data diversity and using multiple training techniques to enhance generalization significantly improves Model 3’s prediction accuracy. (3) Through multi-task training, Model 4 can extract more general residue pair interaction patterns from intra-domain interactions, further strengthening the model’s ability to extract inter-domain residue pair interactions. (4) The recycling mechanism in Model 5 deepens the feature extraction module, further enhancing representation capability. (5) Finally, stepwise fine-tuning allows the language model to update its own parameters to better understand the structural domain features of multi-domain proteins, significantly improving model performance to produce the final DDI_single. The ablation results fully demonstrate that our random sampling, gated cross-attention mechanism, recycling mechanism, multi-task training, stepwise fine-tuning, and various training techniques are all important for DDI_single’s prediction accuracy.

## 6 Conclusion

We presented DDI_single, a single-sequence-based protein domain assembly algorithm that uses a gated cross-attention module to capture inter-domain residue pair interactions. DDI_single achieves *>*20% higher inter-domain distance prediction accuracy than trRosettaX_single and correctly assembles 74.4% of test proteins with known domain conformations (TM-score*>*0.5). When domain conformations are predicted, DDI_single correctly assembles 73.9% of proteins where intra-domain modeling succeeds. The main limitation is dependence on intra-domain prediction quality; future work could integrate DDI_single with stronger single-domain predictors or extend the cross-attention mechanism to simultaneous multi-domain modeling.

## Availability

The core implementation of this work was completed in April 2024; the formal write-up was delayed due to various circumstances. The code and pre-trained models will be made available at https://github.com/Silencezong/DDI_single.

## A Case Studies

### A.1 Assembly with Known Domain Conformations

To intuitively analyze DDI_single’s ability to assemble domains with known conformations, we examine proteins 6icc and 6o5c in detail. Table 5 shows the distance prediction precision, and Table 6 shows the modeling TM-scores.

**Table 5:** Inter-domain distance prediction precision for example proteins with known domain conformations.

| PDB ID | Method | Top10 | Top5 | Top1 |
| --- | --- | --- | --- | --- |
| 6icc | trRosettaX_single | 20% | 20% | 100% |
|  | DDI_single | **40%** | **60%** | 100% |
| 6o5c | trRosettaX_single | 30% | 20% | 100% |
|  | DDI_single | **90%** | **100%** | 100% |

**Table 6:** TM-scores of DDI_single and trRosettaX_single for example proteins with known domain conformations.

| PDB ID | Method | TM-score |
| --- | --- | --- |
| 6icc | trRosettaX_single | 0.6037 |
|  | DDI_single | **0.8359** |
| 6o5c | trRosettaX_single | 0.6006 |
|  | DDI_single | **0.8384** |

DDI_single can accurately identify key inter-residue interactions affecting inter-domain relative positioning, achieving precise domain assembly with overall model TM-scores exceeding 0.7. In contrast, trRosettaX_single’s weaker ability to capture inter-domain interactions means that even when it detects some existing interactions, its failure to identify the key interactions that determine relative positioning results in incorrect domain assembly, with TM-scores around only 0.6. Figure 8 shows the structural superposition for these two examples.

**Figure 8:**
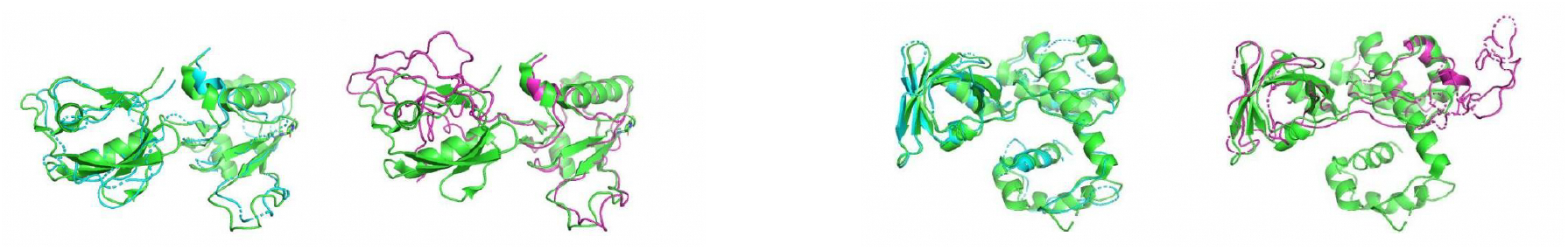
Structural comparison for proteins 6icc (left) and 6o5c (right) with known domain conformations. In each panel, the left superposition shows DDI_single prediction (cyan) vs. native structure (green), and the right superposition shows trRosettaX_single prediction (magenta) vs. native structure (green).

### A.2 Assembly with Unknown Domain Conformations

For proteins 6qmr and 6s73, Table 7 shows inter-domain distance prediction precision and Table 8 shows intra-domain distance prediction precision. Both methods accurately capture inter-domain interactions with very high distance prediction precision. Table 9 shows the modeling TM-scores.

**Table 7:**
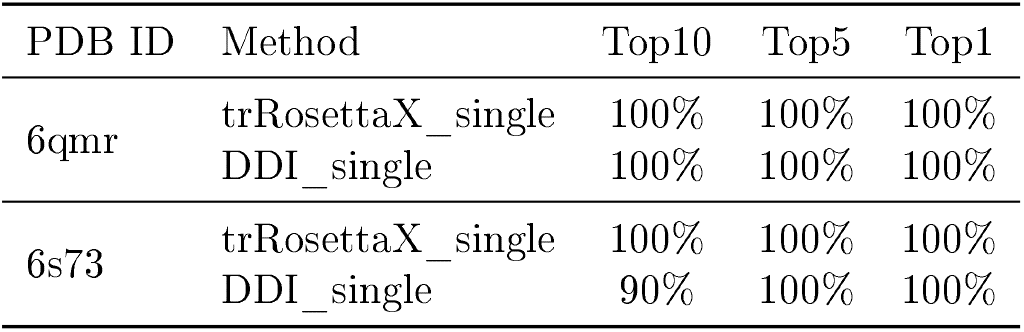
Inter-domain distance prediction precision for example proteins with unknown domain conformations.

| PDB ID | Method | Top10 | Top5 | Top1 |
| --- | --- | --- | --- | --- |
| 6qmr | trRosettaX_single | 100% | 100% | 100% |
|  | DDI_single | 100% | 100% | 100% |
| 6s73 | trRosettaX_single | 100% | 100% | 100% |
|  | DDI_single | 90% | 100% | 100% |

**Table 8:** Intra-domain distance prediction precision by trRosettaX_single for example proteins.

|  | TopL | TopL/2 |
| --- | --- | --- |
| 6qmr | 99% | 100% |
| 6s73 | 93% | 99% |

**Table 9:**
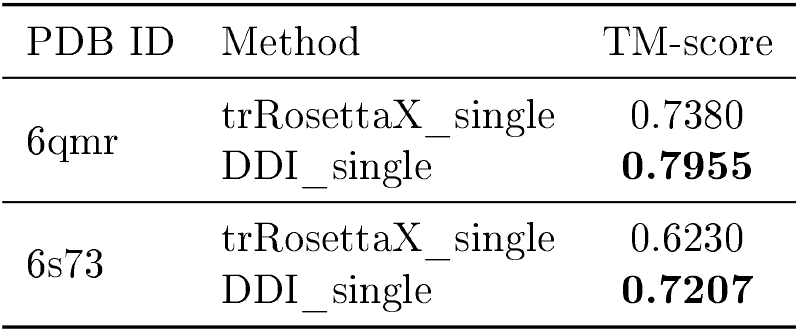
TM-scores of DDI_single and trRosettaX_single for example proteins with unknown domain conformations.

| PDB ID | Method | TM-score |
| --- | --- | --- |
| 6qmr | trRosettaX_single | 0.7380 |
|  | DDI_single | **0.7955** |
| 6s73 | trRosettaX_single | 0.6230 |
|  | DDI_single | **0.7207** |

DDI_single captures the key inter-residue interactions affecting inter-domain relative positioning, successfully achieving accurate domain assembly with overall model TM-scores exceeding 0.7. While trRosettaX_single also captures some inter-domain interactions with similar distance prediction accuracy to DDI_single, it fails to capture the key residue pair interactions that determine inter-domain relative positioning, resulting in the folding algorithm being unable to correctly model the relative positions between domains and producing TM-scores lower than DDI_single. Figure 9 shows the structural superposition for these two examples.

**Figure 9:**
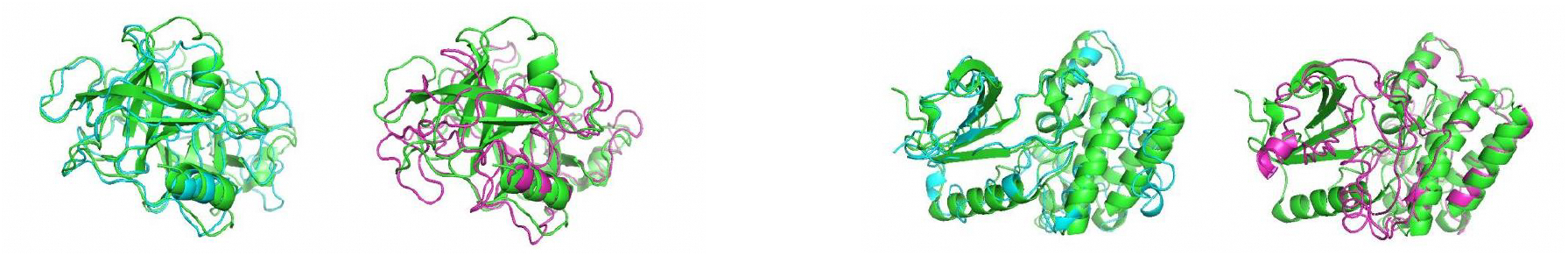
Structural comparison for proteins 6qmr (left) and 6s73 (right) with unknown domain conformations. In each panel, the left superposition shows DDI_single prediction (cyan) vs. native structure (green), and the right superposition shows trRosettaX_single prediction (magenta) vs. native structure (green).

## B Data Preprocessing Details

Multi-domain protein structures are collected from the Protein Data Bank (PDB) [4] using CATH [18] domain annotations. The preprocessing pipeline includes: (1) downloading PDB structures and extracting backbone atom coordinates (*N*, *C*_*α*_, *C*_*β*_, *C*^*′*^); (2) computing inter-residue geometric features for each residue pair—*C*_*β*_—*C*_*β*_ distance *d*, dihedral angle *ω* (rotation of both *C*_*α*_ atoms about the *C*_*β*_—*C*_*β*_ axis), dihedral angle *θ* (rotation of *N*_*i*_ and *C*_*α,j*_ about the *C*_*α,i*_—*C*_*β,i*_ axis), and planar angle *φ* (*C*_*α,i*_—*C*_*β,i*_—*C*_*β,j*_); (3) discretizing *d* into 37 bins (2—20 Å in 0.5 Å steps, plus one bin for *>*20 Å), *ω* and *θ* into 25 bins each (−180° to 180°), and *φ* into 13 bins (0° to 180°); (4) clustering sequences at 40% similarity using MMseqs2 to reduce redundancy; (5) splitting into training/evaluation/test sets by release date.

For glycine residues lacking a *C*_*β*_ atom, a virtual *C*_*β*_ position is computed from backbone geometry. All features are stored as classification labels for cross-entropy training.

## C Training Details

### C.1 Stepwise Fine-Tuning

**Step 1:** Freeze ESM-1b parameters; train only the feature extraction module and distance/orientation prediction module using cross-entropy loss. The model simultaneously predicts both intra-domain and inter-domain distances and orientations, with the inter-domain loss weighted 2× to emphasize inter-domain learning.

**Step 2:** Unfreeze all parameters (including ESM-1b). Add masked language modeling (MLM) [12] as an auxiliary task to prevent catastrophic forgetting of the language model’s learned representations. Continue training with all losses jointly.

### C.2 Training Techniques

- **Optimizer:** Adam [26] with learning rate 10^−5^ and weight decay 10^−4^.
- **Warmup:** Linear warmup [27] over the first 5% of training steps.
- **Dropout:** Rate of 0.3 applied to attention weights and feedforward layers [29].
- **Dynamic masking:** Following RoBERTa [28], masking patterns are regenerated each epoch rather than fixed.
- **Random sampling:** Instead of using a single representative sequence per CATH class, we randomly sample from all sequences in each class per epoch to increase training data diversity.
- **Multi-task training:** Joint prediction of intra-domain and inter-domain distances/orientations, with 2× weight on inter-domain loss to compensate for the smaller number of inter-domain contacts.
- **Batch size:** 1 protein per batch due to variable sequence length; gradients are accumulated over 4 steps.

### D Evaluation Metrics

#### D.1 Precision

Precision measures the accuracy of predicted inter-residue contacts. For each protein, we select the top-*N* residue pairs with highest predicted probability of being in contact (distance 2−8 Å) and compute:

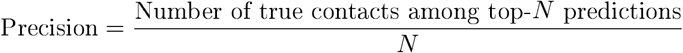

We report Top10, Top5, and Top1 for inter-domain pairs (where *N* = 10, 5, or 1 pairs with highest contact probability), and TopL and TopL/2 for full-length sequences (where *L* is the sequence length).

#### D.2 TM-score

TM-score [30] quantifies the structural similarity between a predicted model and the native structure:

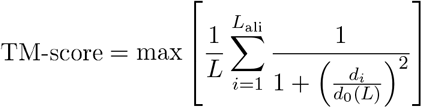

where *L* is the length of the target protein, *L*_ali_ is the number of aligned residues, *d*_*i*_ is the distance between the *i*-th pair of aligned residues after optimal superposition, and 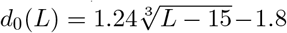 is a length-dependent normalization factor. TM-score ranges from 0 to 1; a score *>*0.5 generally indicates correct fold topology.

